# Active-feedback 3D single-molecule tracking using a fast-responding galvo scanning mirror

**DOI:** 10.1101/2023.03.28.534562

**Authors:** Xiaochen Tan, Shangguo Hou, Chen Zhang, Anastasia Niver, Alexis Johnson, Kevin D. Welsher

**Affiliations:** Duke University; Shenzen Bay Laboratory

## Abstract

Real-time three-dimensional single-particle tracking (RT-3D-SPT) allows continuous detection of individual freely diffusing objects with high spatiotemporal precision by applying closed-loop active feedback in an optical microscope. However, the current tracking speed in RT-3D-SPT is primarily limited by the response time of control actuators, impeding long-term observation of fast diffusive objects such as single molecules. Here, we present an RT-3D-SPT system with improved tracking performance by replacing the XY piezoelectric stage with a galvo scanning mirror with an approximately five-time faster response rate (~5 kHz). Based on the previously developed 3D single-molecule active real-time tracking (3D-SMART), this new implementation with a fast-responding galvo mirror eliminates the mechanical movement of the sample and allows more rapid response to particle motion. The improved tracking performance of the galvo mirror-based implementation is verified through simulation and proof-of-principle experiments. Fluorescent nanoparticles and ~ 1 kB double-stranded DNA molecules were tracked via both the original piezoelectric stage and new galvo mirror implementations. With the new galvo-based implementation, notable increases in tracking duration, localization precision, and the degree to which the objects are locked to the center of the detection volume were observed. These results suggest faster control response elements can expand RT-3D-SPT to a broader range of chemical and biological systems.

## Introduction

Single-molecule spectroscopy (SMS) has been widely used to reveal molecular dynamics obscured by ensemble averaging in chemical and biological systems.^1–4^ In a typical solution-phase SMS measurement, the molecules diffuse extremely rapidly through a focused laser spot, leading to a short observation time on the order of milliseconds. The rapid diffusion of molecules out of the laser focus limits the number of photons and, consequently, the information that can be collected from a single molecule. To extend the observation time, surface tethering (in methods such as wide-field and confocal imaging) or motion confinement (such as Anti-Brownian Electrokinetic trap)^5–7^ have been applied to constrain the molecule’s diffusion and hold it in the detection volume. Surface tethering methods require a potentially perturbative chemical linkage. While trapping methods do not involve a chemical linkage, they still require isolation of the particle from its native environment. A different approach is needed to perform long-duration SMS in native molecular environments.

Real-time three-dimensional single-particle tracking (RT-3D-SPT) eliminates the restrictions mentioned above and allows direct and continuous observation of single, freely diffusing particles in their natural environment, opening access to SMS *in situ*. ^2,8^ RT-3D-SPT uses photons emitted or scattered from a target particle to estimate its position and apply feedback to counteract the particle’s Brownian diffusion, maintaining the particle in the detection volume in real time. The photon information provides spectroscopic data, while the feedback needed to hold the particle in the detection volume yields its 3D trajectory. The temporal resolution of RT-3D-SPT is limited only by the photon detection rate, compared to traditional image-based tracking methods, which are limited by the volumetric imaging rate. This makes RT-3D-SPT a powerful tool to investigate fast single-particle and single-molecule phenomena, such as cellular trafficking,^9,10^ nanoparticle-membrane interactions,^11,12^ DNA transcription,^13–15^ and countless other dynamic processes.^16–18^

Despite the myriad applications described above, there are still limitations to RT-3D-SPT that must be overcome to achieve SMS in all environments truly. The current tracking speed of RT-3D-SPT is barricaded to ~ 20 μm^2^/s or slower with limited tracking duration.^19,20^ As this speed is significantly slower than molecular diffusion in aqueous environments, the addition of high-viscosity solvents is required to perform solution-phase SMS with RT-3D-SPT. Even with increased viscosity, the average tracking duration for single fluorophores is typically below 1 s for most methods.^21–23^ These limitations beg the question: what prevents RT-3D-SPT methods from tracking single molecules in less viscous solutions? The first possibility is that very few photons are emitted by a single molecule during each position update cycle, increasing the uncertainty in position estimation. The limited number of photons becomes further decreased if the molecule diffuses close to the edge of the excitation or detection volume, leading to a higher probability of trajectory termination. The next consideration is the response speed of the control actuator. Suppose the control actuator is responding fast enough. In that case, the probe of interest is more likely to be locked in the center of the detection volume with a correspondingly stable photon detection rate, increasing the trajectory duration. The feedback control of RT-3D-SPT is generally achieved using a piezoelectric stage to move the sample,^19,21,24–26^ or using galvanometric^9,27–30^ or piezoelectric mirrors^10,31,32^ to move the combined excitation/detection volume, leaving the sample stationary. To date, the only methods that have achieved single fluorophore tracking have relied on moving the sample with a piezoelectric stage, despite these stages having a far lower resonant frequency (~1 kHz) compared to galvo scanning mirrors (~10 kHz).^33^ In theory, the faster response element should improve tracking speed, but this has yet to be demonstrated.

One method that has proven particularly successful for real-time single-molecule tracking is 3D single-molecule active real-time tracking (3D-SMART). 3D-SMART uses a rapidly moving laser excitation spot over a relatively large range (1 μm × 1 μm × 2 μm) to reduce the likelihood of a molecule leaving the detection range.^14,20^ As a result, 3D-SMART can track single ATTO 647N fluorophores in 90% glycerol solution with an average tracking duration of more than 15 s.^14^ Previously demonstrated 3D-SMART results for tracking dsDNA provide an interesting test case for the factors that limit trajectory duration in RT-3D-SMT. Interestingly, with the same labeling condition (one fluorophore per strand), 3D-SMART can track single 99 bp dsDNA in 90% glycerol for ~17 s on average, while the tracking duration drops to ~ 1 s in 40% glycerol.^14^ These data suggest that trajectory termination is not simply due to insufficient photon budget but rather is limited by the physical response time of the control actuator, in this case, a piezoelectric stage. The response time of the stage is ~ 1 ms in all three dimensions, making it impossible to follow a single small molecule in water. Faster control might be achieved using more aggressive control parameters, but this will come with increased overshoot and oscillation. In this paper, we demonstrate that the response time in RT-3D-SPT can indeed be dramatically improved by implementing a galvo scanning mirror into 3D-SMART. Instead of moving the sample to compensate for the particle’s motion, the galvo scanning mirror deflects the laser beam to follow the particle with a response time that is five-fold faster than the previous piezoelectric stage (~210 μs, Fig. 1b, c). Tracking feedback parameters were optimized, and proof-of-principle experiments were performed to evaluate the lateral tracking performance of 3D-SMART with fluorescent nanoparticles and dsDNA. The preliminary results show that the galvo scanning mirror implementation improves the response time in RT-3D-SPT and that highspeed steering mirrors provide a path towards active-feedback single-molecule tracking in all environments.

**Figure 1.**
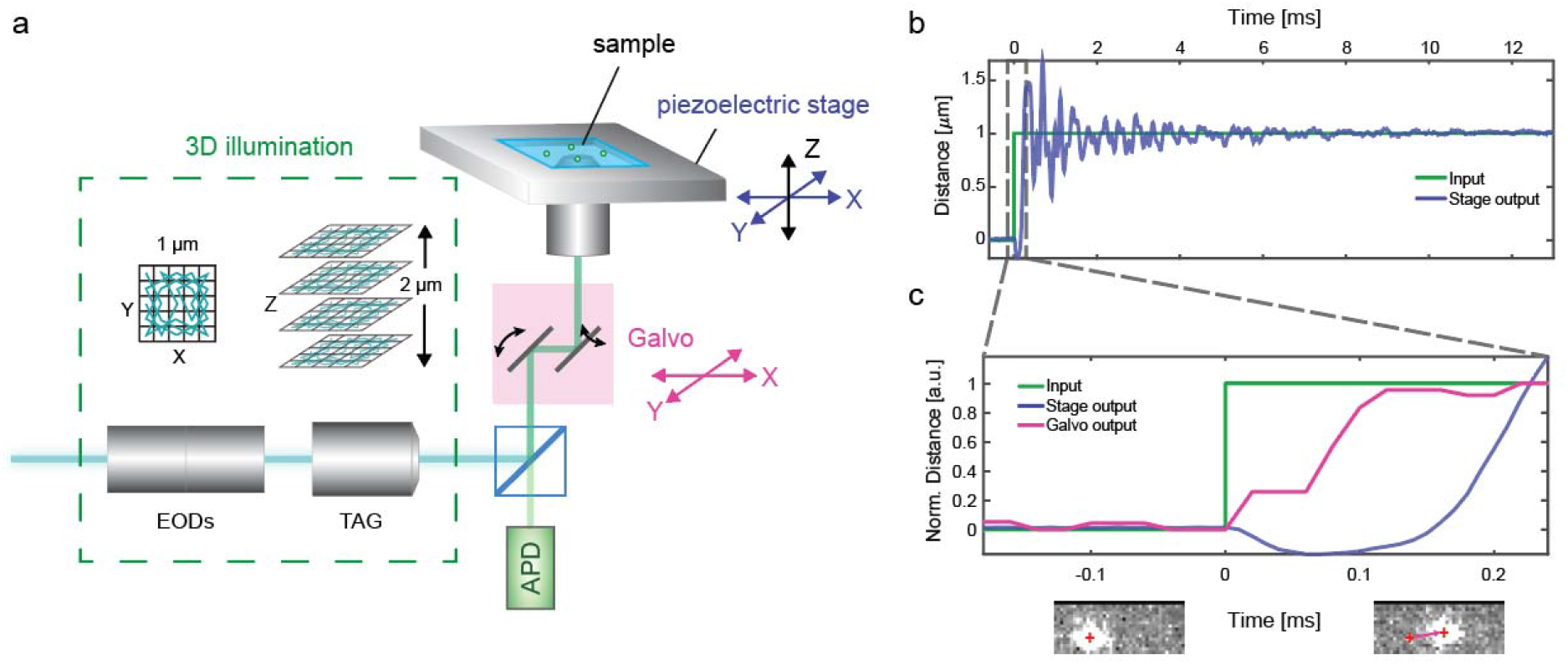
Experimental setup. (a) Schematic illustration of the 2D galvo mirror implemented into the 3DSMART setup. (b) Piezoelectric stage (XY) step response (blue) to a step input (green), measured by direct readout. (c) Galvo step response (pink) and zoomed stage response (blue) to a step input (green). Images show the centroid position (red cross) of the laser on the sCMOS camera before and after the step input.

## Experimental section

### Experimental setup

The setup is based on the previously developed 3D-SMART, with the addition of a 2D galvo scanning mirror for XY tracking (Fig. 1a, Fig. S1). The 3D-SMART microscope utilizes a 2D electro-optical deflector (EOD, M310A, ConOptics) to deflect the laser beam in the lateral direction in a 5 × 5 Knight’s Tour (KT) pattern,^34^ which uniformly samples the scan area with minimal lag compared to a simple raster scan. The range and number of pixels in the KT are adjustable based on the experimental needs. Here, the scanning range is set to 1 μm × 1 μm with a dwell time of 20 μs per spot, the same as the original 3D-SMART implementation. For axial excitation, the laser is then sent through a tunable acoustic gradient (TAG, TAG 2.0, TAG Optics) lens to deflect the laser beam in a sinusoidal pattern at a frequency of 70 kHz.^35^ The amplitude is set to 35% of the maximum, corresponding to a ~2 μm axial range. Combined, a rapid illumination pattern over a 1 μm × 1 μm × 2 μm range is generated to excite the fluorescent object of interest.

Once the fluorescent object diffuses into the 3D excitation volume, the emitted photon arrivals are collected by a single-photon counting avalanche photodiode (APD, SPCM-ARQH, Excelitas). The photon arrivals and laser positions are fed into an assumed Gaussian density Kalman filter to calculate the particle’s position relative to the 3D scan center in real time.^36^ The original implementation of 3D-SMART uses a piezoelectric nanopositioner (Nano-PDQ275HS, Mad City Labs) to move the sample in XY and a piezoelectric lens positioner (Nano-OP65HS, Mad City Labs) to move the objective lens in Z to maintain the tracked object in the center of the 3D scan volume. The response time of the piezoelectric stage is measured by directly measuring the readout signal following a command, yielding a ~ 1 ms response time. When overshoot and oscillation are taken into account, the overall settling time is ~ 5 ms (Fig. 1b, Fig. S2). To this setup, a 2D galvo scanning mirror system (SG7220-A, SINO Galvo) is added for XY tracking as a direct comparison to the prior stage-based XY tracking. The 2D galvo mirror scans the tracking excitation spot, and the resulting fluorescence signal is de-scanned and collected by the APD. The EODs, TAG lens, piezoelectric stage, galvo mirror, and single-photon counting APDs are fed into a field-programmable gate array (FPGA, NI-7852r, National Instruments). The response time of the galvo mirror was measured by recording the reflected laser spot position with a high framerate sCMOS camera (pco.edge 4.2, PCO) in a very small region of interest (ROI). Figure 1c shows the measured step response by averaging three sCMOS movies (frame time = 76 ± 13 μs). The centroid of the laser position (as indicated by the red mark in Fig. 1c) is identified in each frame and used to calculate the normalized distance traveled by the laser spot. Overshoot and oscillation of the galvo are not observed due to the limited frame rate of the camera.

### Position estimation and feedback

Photon arrivals recorded by the APD are tagged with XYZ position (based on EOD and TAG command positions) and fed into the Kalman filter to estimate the current particle position. The position estimation algorithm is verified by scanning an immobilized nanoparticle through the tracking excitation volume using the piezoelectric stage (Fig. S3). Once the updated position estimate is acquired, an integral controller is used to move the piezoelectric stage (for the original 3D-SMART) or the galvo mirror to bring the particle back to the center of the scanning volume. More details on the position estimation algorithm and feedback controller can be found in the SI.

### System calibration

The size of mirrors and lenses in the system is the main factor that affects the tracking range of the galvo mirror-based system. If the mirror deflection angle is too large, the laser beam will be clipped by the mirror or lens aperture, affecting the system point spread function (PSF). The galvo scanning range was measured using a slide with fluorophores uniformly distributed on the surface to avoid this vignetting. Figure S4 shows that a uniform intensity is collected over a ~51 μm scanning range of the galvo mirror. The position of the objective lens, which is moved to hold the particle in focus, can also affect the excitation laser power as more or less of the scanned beam is passed through the back aperture of the lens. The correction for this excitation power fluctuation is shown in Figure S4 and can be mitigated using real-time laser power modulation if needed. The galvo voltage-distance conversion was measured by moving an immobilized nanoparticle using the piezoelectric stage and tracking it using the galvo mirror (Fig. S5).

## Results and Discussion

### Real-time 3D single-particle tracking using a 2D galvo mirror

The 2D tracking performance of the galvo and stage were simulated to validate the effectiveness of the galvo implementation. The simulation was performed based on the impulse responses of the stage and galvo, attained by fitting the measured step responses (Fig. 1b, c). From the simulation results, galvo tracking exhibited longer trajectory duration when tracking particles with diffusion coefficients greater than 20 μm^2^/s given sufficient photons (Fig. S6).

The experimental tracking performance was evaluated using 110 nm diameter fluorescent polystyrene nanoparticles (NPs). Figure 2 shows example trajectories of both stage and galvo tracking. The 3D trajectories were recorded based on the stage and galvo motion (Fig. 2a). The intensity trace and photon arrivals were simultaneously acquired during each trajectory (Fig. 2b). Mean square displacement (MSD, see SI) analysis shows the representative trajectories are particles of similar size (Fig. 2c). To quantitatively evaluate tracking performance, the “central photon fraction” was calculated from the photon arrivals. The central photon fraction is the number of photons that are collected from the center pixel in the 25-point KT scan, divided by the total number of photons collected (as indicated by the green box in Fig. 2b). This parameter reflects how tightly the particle is locked in the center of the scan area while being actively tracked. As a reference, immobilized NPs have a central photon fraction of 0.091 ± 0.002 (n = 24, Fig. S7). For the representative freely diffusing particle trajectories shown in Figure 1, the central photon fraction was calculated to be 0.083 and 0.085 for stage and galvo tracking, respectively.

**Figure 2.**
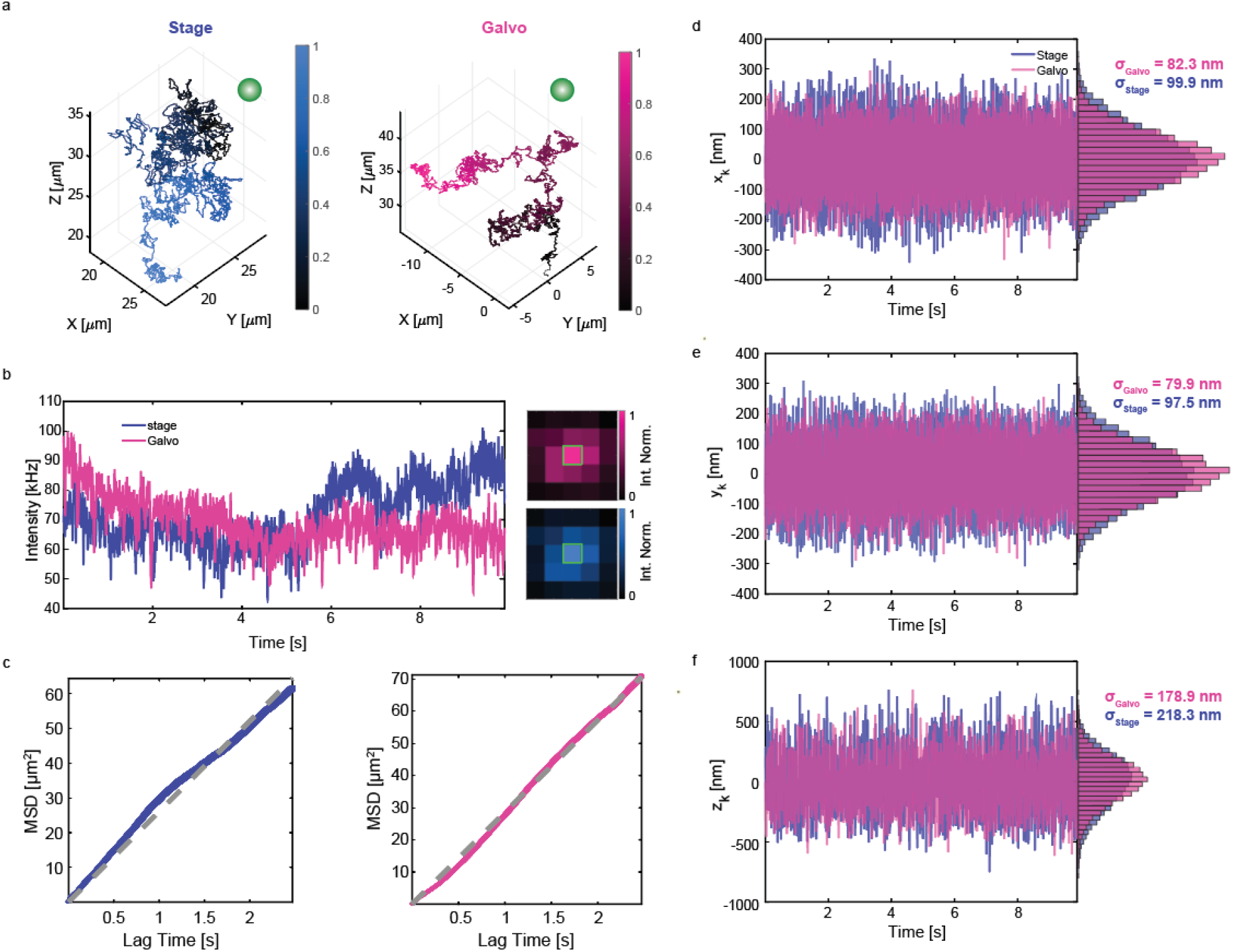
Example stage and galvo single-particle trajectories. (a) Example trajectories. Blue: stage tracking. Pink: galvo tracking. The galvo trajectory duration is cropped to match the stage tracking duration (t = 9.84 s) for ease of comparison. The color bar indicates the normalized tracking duration. (b) Intensity traces and photon arrivals. The green box indicates the central pixel. (c) MSD measurements. The grey dotted line indicates the linear fit. D = 4.37 μm^2^/s for stage tracking, D = 4.75 μm^2^/s for galvo tracking. (d-f) X, Y, and Z position estimates during stage (blue) and galvo (pink) tracking.

The estimated positions relative to the scan center (x_k_, y_k_, and z_k_) are recorded in realtime during each trajectory to evaluate particle displacement from the center of the scan area. The distribution of these estimates shows that the galvo tracking method holds the NP more tightly locked while tracking (Fig. 2d-f). There is also a noticeable improvement in the Z localization compared to the stage tracking case, despite both methods using the same piezoelectric nanopositioner for Z tracking. This is likely a result of the improved XY tracking of the galvo method.

### Feedback control parameter optimization

Since the galvo and stage have different response times (Fig. 1b, c), each method’s integral control parameters must be optimized for a fair comparison. Tracking performance on 110 nm diameter NPs (average intensity 174 ± 81 kHz, Fig. S8) was evaluated for different integral control parameter (K_I_) values (0.0025 to 0.125, Fig.3, n > 30 for each K_I_). The effect of the integral control parameter was quantified using the following three parameters: trajectory duration (Fig. 3a), localization precision (Fig. 3b,d), and central photon fraction (Fig. 3c). All NP aggregates, trajectories shorter than 0.1 s in duration, or that ended due to reaching the limit of the stage or galvo motion were removed from further analysis (See Fig. S9 for the end positions of all trajectories). The maximum tracking duration threshold was set to 20 s (grey dashed line in Fig. 3a) to avoid the effects of photobleaching. There is a striking increase in the trajectory duration for galvo tracking as compared to stage tracking for all K_I_ values without exception (Fig. 3a). Furthermore, the average galvo tracking duration hits the maximum set threshold of 20 s for any K_I_ value between 0.0325 and 0.0925, indicating a more consistent tracking performance. The standard deviation of the particle’s position estimate within the KT yields the localization precision (Fig. 3b,d). The X and Y precision were optimized when K_I_ = ~0.0425 for galvo tracking and K_I_ = ~0.0225 for stage tracking. Given a feedback loop time of 20 μs, these correspond to response times of 0.471 ms and 0.889 ms, respectively, reflecting the improved response speed of the galvo mirror. A similar trend is observed in the central photon fraction plots, which show a broad range of K_I_ values resulting in nearly identical performance for galvo tracking. These results show that the range of acceptable K_I_ values is far larger for galvo tracking compared to stage tracking. These data are again consistent with the idea that the response of the galvo mirror is faster than the stage, making more aggressive control parameters viable. The same experiments were also conducted under lower laser excitation power to evaluate low photon count rate conditions (average NP intensity below 12 kHz, Figs. S8 and S10). For dimmer particles, the tracking effectiveness is more likely to be determined by the lack of photons and the corresponding decreased signal-to-background ratio. As a result, the dimmer NPs show narrower peaks of optimal K_I_ values compared to brighter NPs. However, even under these more difficult tracking conditions, the galvo implementation still exhibits better peak performance than stage tracking. Simulated trajectories showed similar trends, showing a wide range of acceptable K_I_ values and reduced tracking error for galvo tracking compared to stage tracking (Fig. S11).

**Figure 3.**
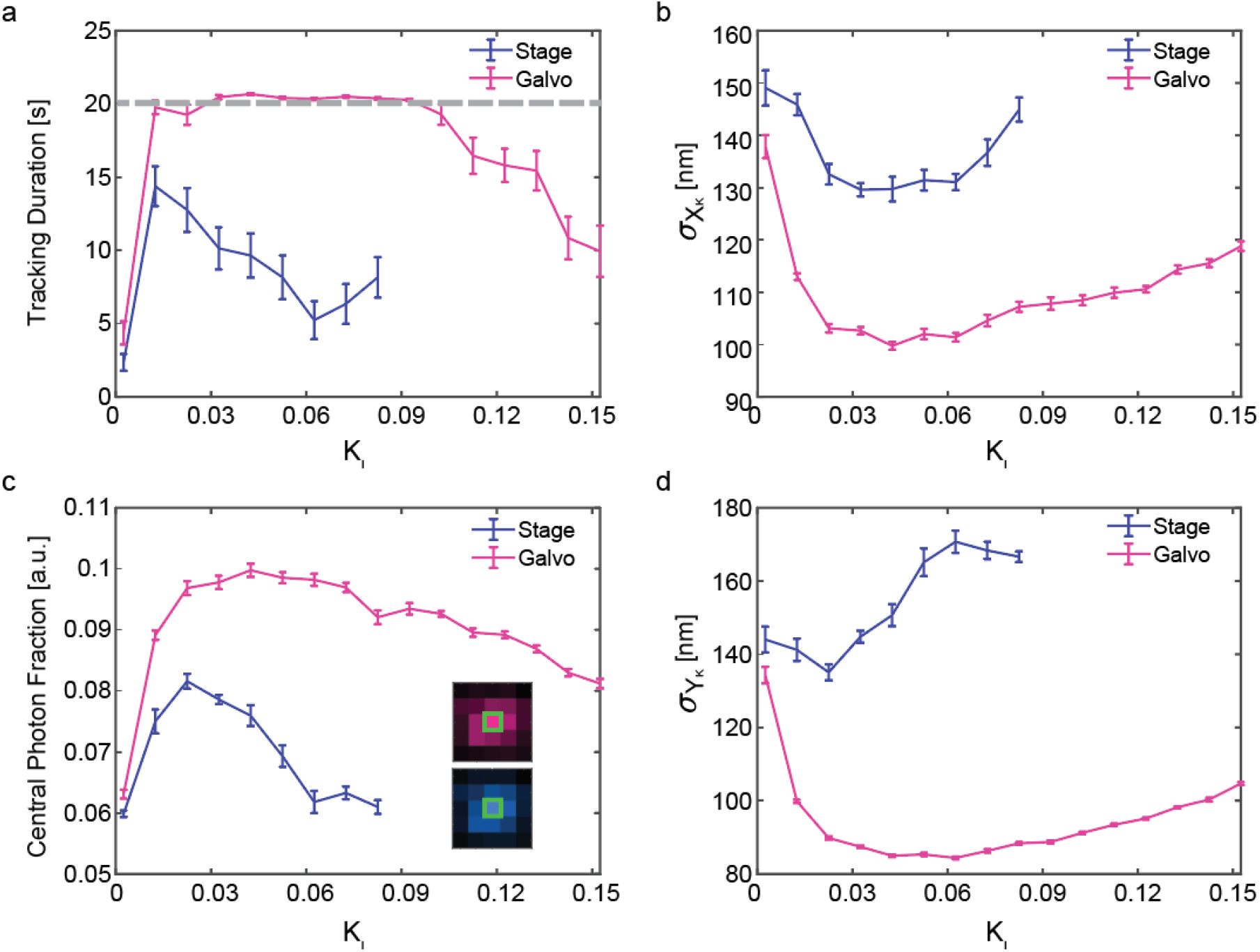
Tracking performance as a function of integral feedback control (K_I_) values. (a) Average tracking duration. The grey dashed line is the set tracking duration threshold. (b) X precision averaged over n > 30 trajectories. (c) Average central photon fraction. (d) Y precision averaged over n > 30 trajectories. The error bars indicate the standard error.

### Tracking with an information-efficient scan pattern

Recent work by Zhang et al. has shown that while the KT scan works well for high-speed tracking, a 4-point scanning pattern at the edges of a well-centered particle leads to increased localization precision.^37^ The problem with applying such a minimalist scan is that while a well-centered particle is localized with more precision, rapidly diffusing particles can quickly move to the extremes of the grid and be lost. Given the success of the galvo approach above to hold the diffusing particle closer to the scan center, it is hypothesized here that the galvo approach would increase tracking duration for the information-efficient 4-point scanning pattern.

To test this hypothesis, stage versus galvo tracking performance was compared with the same integral control parameter of 0.0125. This parameter value is selected to be consistent with the value applied in the previous 3D-SMART work that, while not optimized, works well for both stage and galvo tracking. Galvo and stage tracking were performed with the original 5×5 KT, the information-efficient 4-point scan (Fig. 4), as well as a smaller 3×3 KT (Fig. S12). Figure 4b shows the histograms of tracking duration of the stage and galvo tracking with the 5×5 KT. Consistent with the results above, the galvo shows increased tracking duration (75.7 ± 52.8 s) as compared to stage tracking (58.2 ± 56.0 s, Fig. 4b, p < 0.05, two-tailed t-test). The increased stability of the galvo also translates to the 4-point scan (galvo = 54.1 ± 56.2 s, stage = 30.5 ± 51.5 s, p< 0.01). As expected from the prior work, the localization precision of the 4-point scan is dramatically improved compared to the 5×5 KT (Fig. 4c, d). Notably, the galvo can achieve this increased precision while matching the tracking duration of the stage for the 5×5 KT.

**Figure 4.**
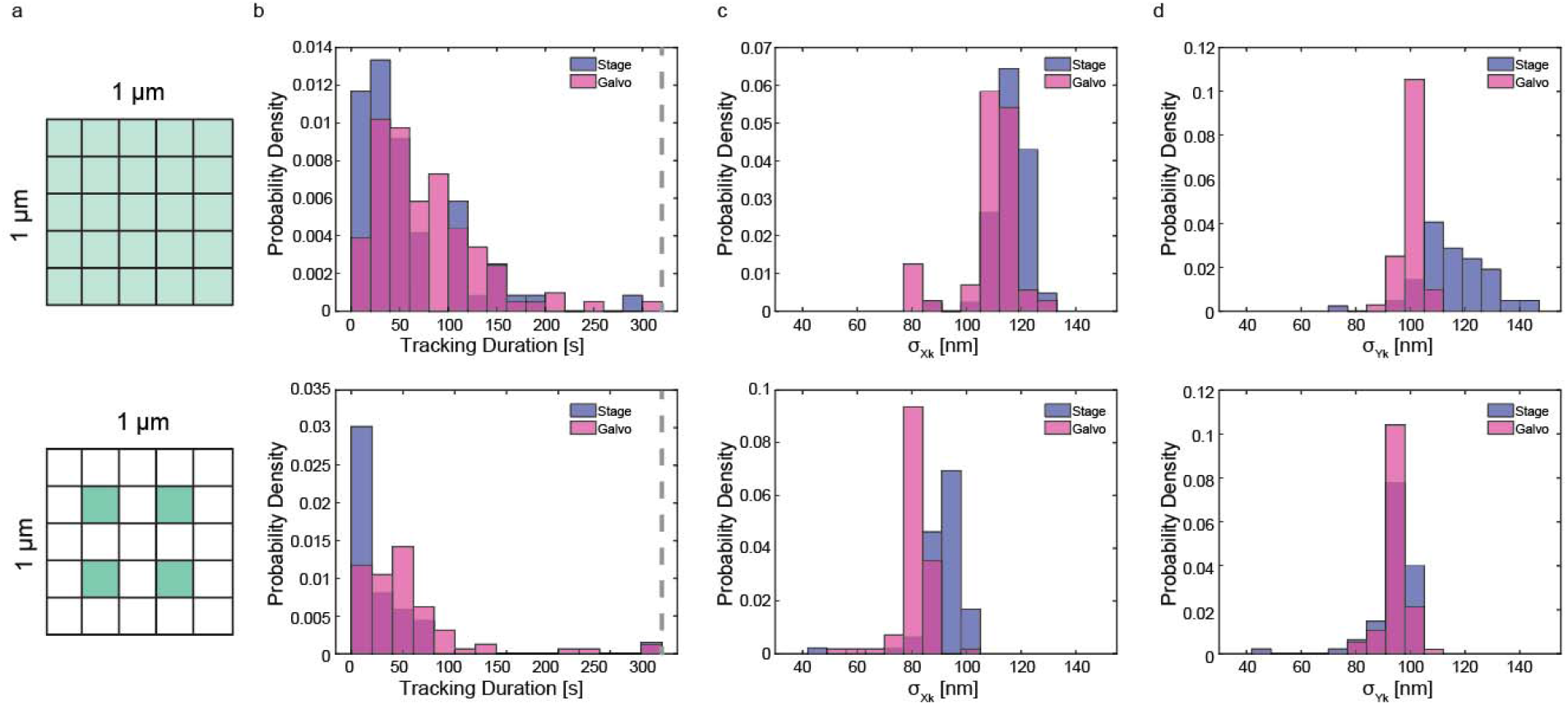
Tracking with an information-efficient 4-point scan pattern. (a) 5×5 KT (top) and information-efficient 4-point scan (bottom) laser scanning patterns. Pixel size = 200 nm. Green pixels are the pixels scanned by the tracking excitation laser. (b) Histograms of galvo (pink) and stage (blue) tracking durations. Top: Tracking durations are 58.2 ± 56.0 s for stage (n = 60) and 75.7 ± 52.8 s for galvo (n = 103, p < 0.05, two-tailed t-test). Bottom: Tracking duration is 30.5 ± 51.5 s for stage (n = 68) and 54.1 ± 56.2 s for galvo (n = 81, p < 0.01). The grey dashed lines are the tracking duration threshold. (c) Histograms of localization precision in X. Top: 116.4 ± 6.5 nm for stage, 108.8 ± 10.6 nm for galvo (p < 0.01). Bottom: 91.1 ± 8.3 nm for stage, 81.5 ± 5.8 nm for galvo (p < 0.01). (d) Histograms of localization precision in Y. Top: 115.2 ± 12.6 nm for stage and 100.9 ± 4.0 nm for galvo (p < 0.01). Bottom: 93.7 ± 8.2 nm for stage, 94.8 ± 4.1 for galvo (p > 0.05). Data are expressed as mean ± standard deviation.

### Active-feedback 3D single-molecule tracking

Given the data above, the faster responding galvo scanning mirrors are expected to improve the speed, precision, and duration in real-time 3D single-molecule tracking. As a demonstration, single-molecule tracking experiments were conducted on YOYO-1 labeled dsDNA in water (1136 bp, Fig. 5a,b). Tracked YOYO1-dsDNA molecules exhibit an average diffusion coefficient of 7.4 ± 3.3 μm^2^/s and a corresponding radius of 34.2 ± 18.0 nm, showing good agreement with prior experiments and the theoretical wormlike chain (WLC) model (green dashed line in Fig. 5c).^14,38,39^ Compared to stage tracking, an increase in tracking duration (from 6.4 ± 5.1s to 8.0 ± 4.9 s, p < 0.01) and central photon fraction (from 0.070 ± 0.004 to 0.077 ± 0.005, p < 0.01) is revealed for galvo tracking (Fig. 5d). While the tracking duration is primarily limited by photobleaching (Fig. 5b), the central photon fraction shows a dramatic improvement for the galvo approach (Fig. 5d), indicating more precise single molecule tracking.

**Figure 5.**
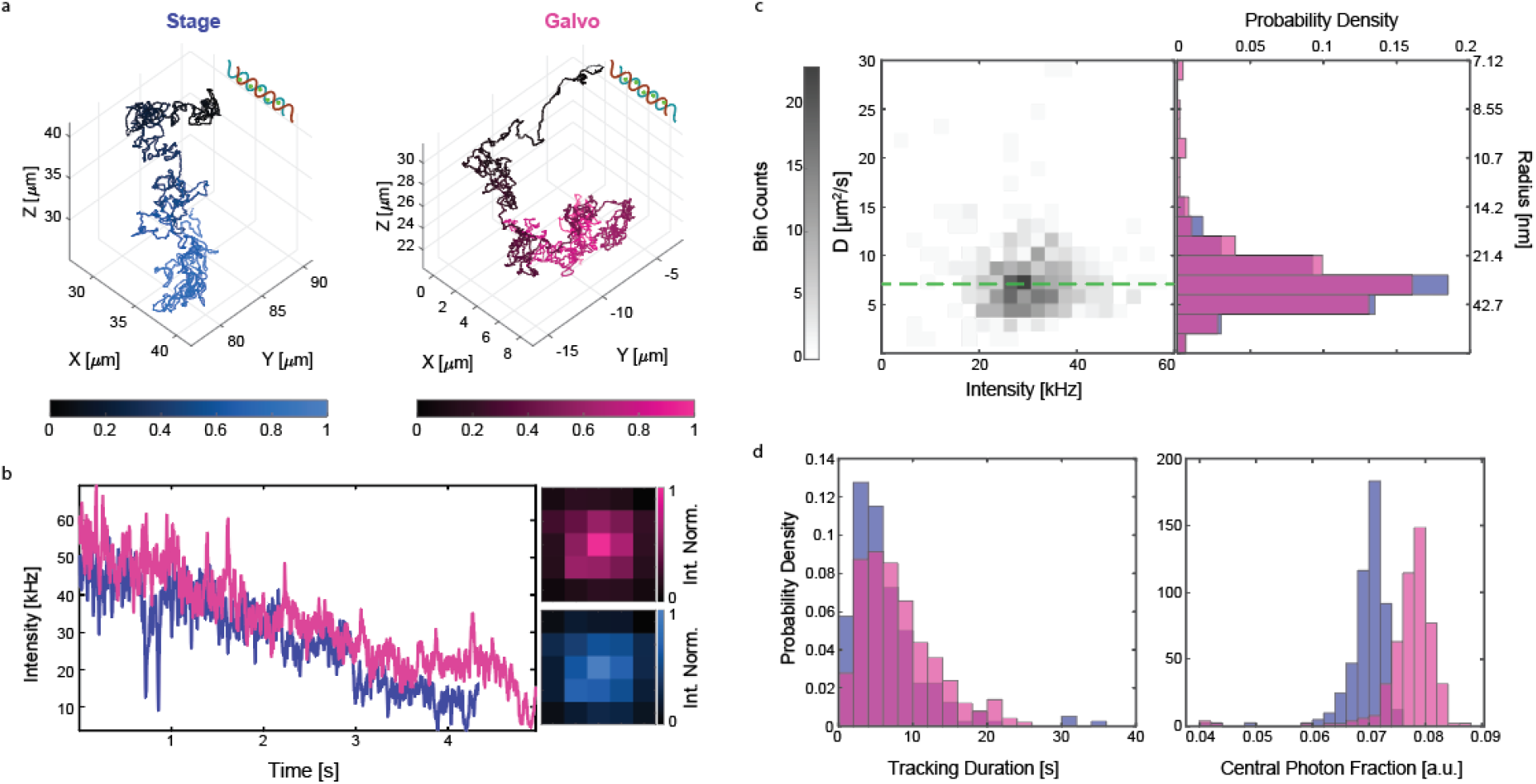
YOYO-1 labeled dsDNA (1132 bp) tracking. (a) Example trajectories, (b) intensity traces and photon arrival distributions for stage and galvo tracking of single YOYO-1 labeled dsDNA at K_I_ = 0.0125. (c) 2D histogram of diffusion coefficient versus intensity for single dsDNA trajectories. The green line indicates the theoretical diffusion coefficient based on the WLC model. (d) Histogram of tracking duration. Stage tracking (blue): 6.4 ± 5.1 s, n = 200; galvo tracking (pink): 8.0 ± 4.9 s, n = 252 (p < 0.01). (e) Histogram of central photon fraction. Stage tracking (blue): 0.070 ±0.004; galvo tracking (pink): 0.077 ± 0.005 (p < 0.01). Data are expressed as mean ± standard deviation.

## Conclusion

This work demonstrates that applying feedback with a high-speed galvo mirror improves the performance in real-time 3D single-particle tracking. The galvo implementation showed greater trajectory duration and localization precision when tracking fluorescent NPs. In addition, the faster feedback response held freely diffusing particles closer to the scan center, enabling the application of an information-efficient 4-point scan that led to dramatically improved localization precision. Finally, galvo-based tracking showed increased precision in tracking single dsDNA in water.

Overall, these results support the central thesis of this work that the piezoelectric stage feedback control element is a limiting factor in real-time 3D single-particle tracking. While the results presented herein are promising, they are still limited by the relatively slow response of the piezoelectric lens positioner for Z tracking. The challenge in future work will be to extend this galvo control approach to three dimensions. Commercially available 3D galvo scanners typically use a motorized lens positioner for focus control, which will have speed comparable to or worse than the piezo lens positioner used in this work. More innovative approaches, such as recently developed galvo-driven focus systems^40^ or deformable mirrors^41,42^ could provide a means to a faster response along the axial dimension. In addition to the improved tracking performance, a full 3D galvo implementation would be able to perform active-feedback 3D single-molecule tracking with a fully stationary sample, which is critical for interfacing with complementary imaging methods and translation of this powerful method to more complex biological and chemical systems.

