## Supplementary Information for "Active-feedback 3D single-molecule tracking using a fast-responding galvo scanning mirror"

### Supplementary Methods

#### Particle position estimation

The photon arrivals (recorded from APD) and the associated laser position (recorded from EOD and TAG lens command positions) are fed into a Kalman filter with the following assumptions: (i) there is zero covariance between X, Y, and Z dimensions; (ii) the likelihood distribution is normally distributed; (iii) the signal is much larger than the background; (iv) there is less than one photon count per bin on average. The particle's position prediction is given based on the assumption of Brownian diffusion:

$$\hat{x}_{k|k-1} = \hat{x}_{k-1|k-1} \tag{1}$$

$$\hat{\sigma}^2_{k|k-1} = \hat{\sigma}^2_{k-1|k-1} + 2D\tau \tag{2}$$

The position estimation is updated by:

$$\hat{x}_{k|k} = \frac{\hat{x}_{k|k-1}\omega^2 + c_k n_k \hat{\sigma}^2_{k|k-1}}{\omega^2 + n_k \hat{\sigma}^2_{k|k-1}} \tag{3}$$

$$\hat{\sigma}^2_{k|k} = \frac{\hat{\sigma}^2_{k|k-1}\omega^2}{\omega^2 + n_k \hat{\sigma}^2_{k|k-1}} \tag{4}$$

Here the $\hat{x}_{k|k}$ is the particle's estimated position, $\hat{x}_{k|k-1}$ is the previous position estimate, $\omega^2$ is the laser beam covariance, $c_k$ is the laser spot position, $n_k$ is the number of photons collected. $\hat{\sigma}^2_{k|k-1}$ is the variance of the previous position estimate, $\hat{\sigma}^2_{k|k}$ is the variance of the current position estimate. Position estimation is verified by scanning an immobilized nanoparticle (NP) with EOD and TAG lens using a piezoelectric stage (Fig. S3).

#### Galvo mirror/piezoelectric stage control

Once the updated position estimate is acquired, a control signal is fed to the piezoelectric stage (for the original 3D-SMART) or the galvo mirror to bring the particle back to the center of the scanning volume. The stage/galvo movement command ($x_{com}$) at each step is determined by an integral controller:

$$x_{com} = K_I \sum x_{k|k} \tag{5}$$

Where $K_I$ is the integral gain. The actual stage/galvo motion ($\mathrm{x_{act}}$) is the convolution of the command ($\mathrm{x_{com}}$) and the system impulse function (h):

$$x_{act}(t) = x_{com}(t) * h(t) \tag{6}$$

The impulse response functions of stage and galvo were derived from the fit of the measured step responses using MATLAB system identification toolbox.

#### MSD calculations

The particle's 3D coordinates (x, y, and z) are used for the mean square displacement (MSD) measurement:

$$\mathrm{MSD}(\Delta t) = \frac{1}{N}\sum_{i=1}^{N}[(x(i+\Delta t) - x(i))^2 + (y(i+\Delta t) - y(i))^2 + (z(i+\Delta t) - z(i))^2] \quad (7)$$

where N is the number of data points (N), and Δt is the lag time. The diffusion coefficient (D) is obtained by fitting the MSD versus the lag time based on Brownian diffusion assumption. Next, the radius can be calculated using the Stokes-Einstein equation:

$$D = \frac{RT}{N_A}\frac{1}{6\pi\eta r} \quad (8)$$

where R is gas constant, $N_A$ is Avogadro's number, η is the dynamic viscosity of the solution, and r is the hydrodynamic radius of the tracked particle.

**Materials and sample preparation**

The dye slide used for calibration was purchased from Ted Pella, Inc. The 110 nm Dragon green polystyrene nanoparticles were purchased from Bangs Laboratories, Inc. Immobilized beads were prepared by diluting the NPs in PBS (1X) buffer to ~1 pM concentration. Freely diffusing beads were diluted in deionized $H_2O$ to ~10 pM for tracking.

**DNA preparation and labeling**

dsDNA molecules (1136 bp) were prepared by PCR (BioRad) using the following primers:

Forward primer: CGCAAATGGGCGGTAGGCGTG

Reverse primer: TTTTCCTGCAGGGCTCTCAAGCGCGG

The following sequence was used:

CGCAAATGGGCGGTAGGCGTGtacggtgggaggtctatataagcagagctctctggctaactagaga
acccactgcttactggcttatcgaaattaatacgactcactatagggagacccaagctggctagcgtttaaacttaagct
tcgaattctgcagtcgacgAccggtcgccaccatggtagcaggtcatgcctctggcagccccgcattcgggaccgcc
tctcattcgaattgcgaacatgaagagatccacctcgccggctcgatccagccgcatggcgcgcttctggtcgtcagc
gaacatgatcatcgcgtcatccaggccagcgccaacgccgcggaatttctgaatctcggaagcgtactcggcgttcc
gctcgccgagatcgacggcgatctgttgatcaagatcctgccgcatctcgatcccaccgccgaaggcatgccggtcg
cggtgcgctgccggatcggcaatccctctacggagtactgcggtctgatgcatcggcctccggaaggcgggctgatc
atcgaactcgaacgtgccggcccgtcgatcgatctgtcaggcacgctggcgccggcgctggagcggatccgcacg
gcgggttcactgcgcgcgctgtgcgatgacaccgtgctgctgtttcagcagtgcaccggctacgaccgggtgatggtg
tatcgtttcgatgagcaaggccacggcctggtattctccgagtgccatgtgcctgggctcgaatcctatttcggcaaccg
ctatccgtcgtcgactgtcccgcagatggcgcggcagctgtacgtgcggcagcgcgtccgcgtgctggtcgacgtca
cctatcagccggtgccgctggagccgcggctgtcgccgctgaccgggcgcgatctcgacatgtcgggctgcttcctgc
gctcgatgtcgccgtgccatctgcagttcctgaaggacatgggcgtgcgcgccaccctggcggtgtcgctggtggtcg
gcggcaagctgtggggcctggttgtctgtcaccattatctgccgcgcttcatccgtttcgagctgcgggcgatctgcaaa
cggctcgccgaaaggatcgcgacgcggatcaccgcgcttgagagcCCTGCAGGAAAA

The concentration of stock DNA (134ng/μL) was measured by a spectrophotometer (BioSpec-nano, SHIMADZU). The DNA molecules were fluorescently labeled by YOYO-1 dyes (ThermoFisher Scientific) for tracking experiments. 1.2 μL of stock DNA solution was added to 400 μL of 10,000X of stock dye solution in TAE buffer. The mixture was incubated for 60 min at room temperature while shaking, then centrifuged using a 30 kDa spin filter to remove the unbound dyes to reduce the background signal. The mixture was resuspended in PBS buffer for monthly storage. The purified solution was diluted 50 times in deionized $H_2O$ (to prevent the DNA from precipitating to the coverslip) for tracking experiments.

### Supplementary Figures

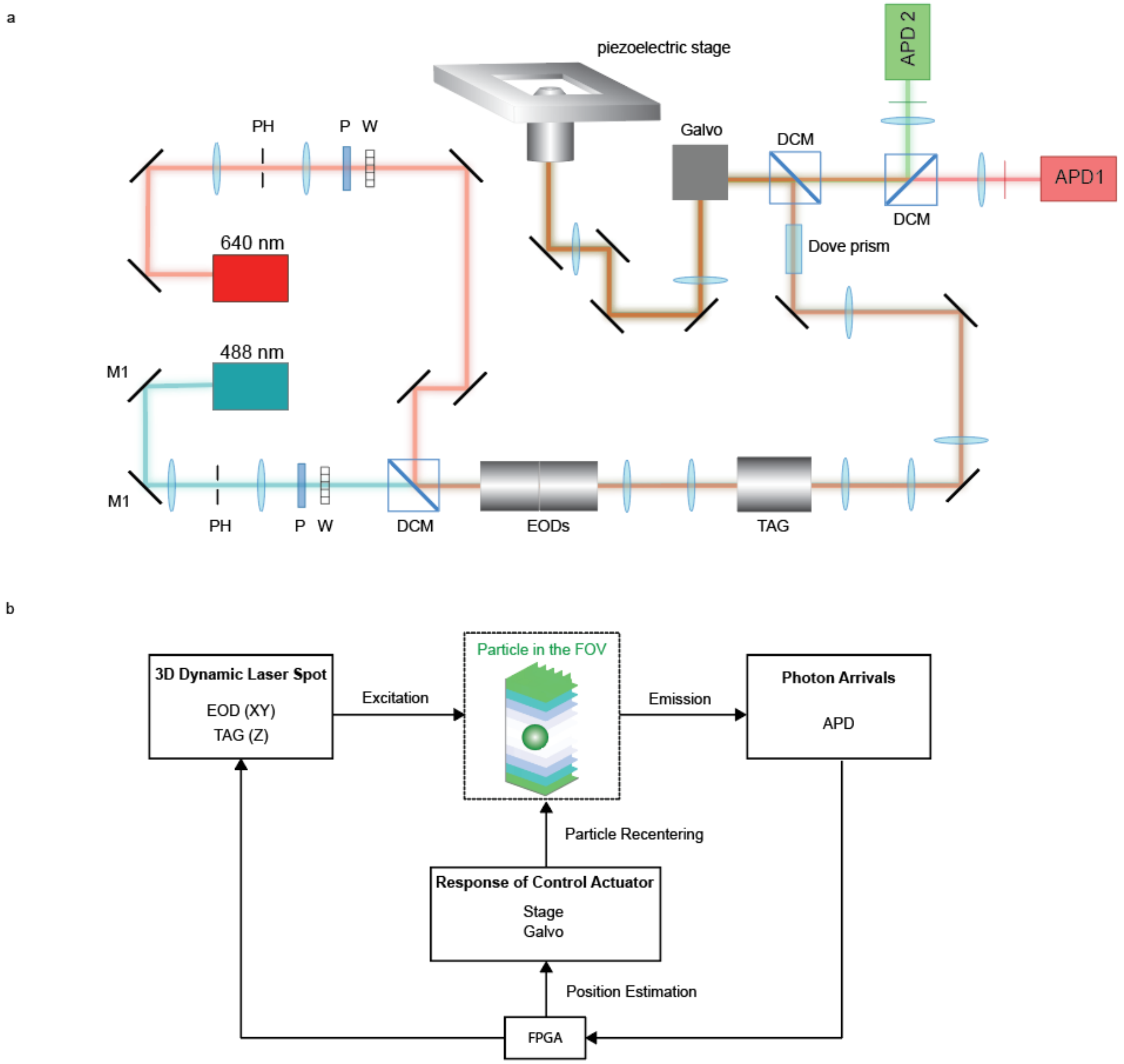

**Figure S1. Real-time 3D single-particle tracking with galvo scanning mirror and piezoelectric stage.** (a) Optical setup. M: mirror. PH: pinhole. P: Glan-Thompson polarizer. W: half-wave plate. DCM: dichroic mirror. EODs: electro-optical deflectors. TAG: tunable acoustic gradient lens. F: emission filter. APD: avalanched photodiode. (b) Tracking principle. FPGA: field-programmable gate array.

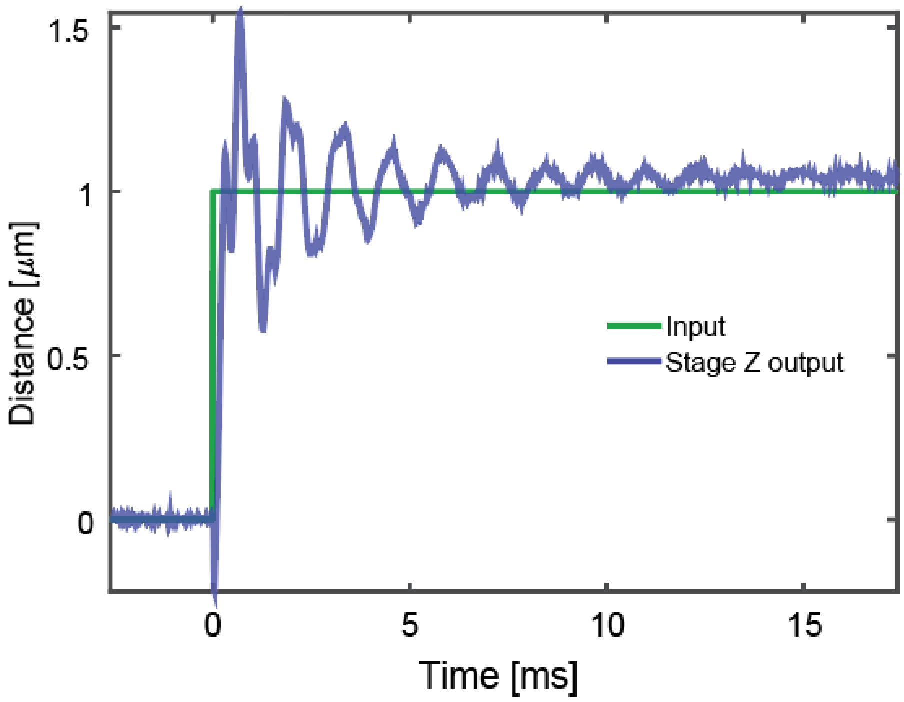

**Figure S2. Piezoelectric stage step response in Z.**

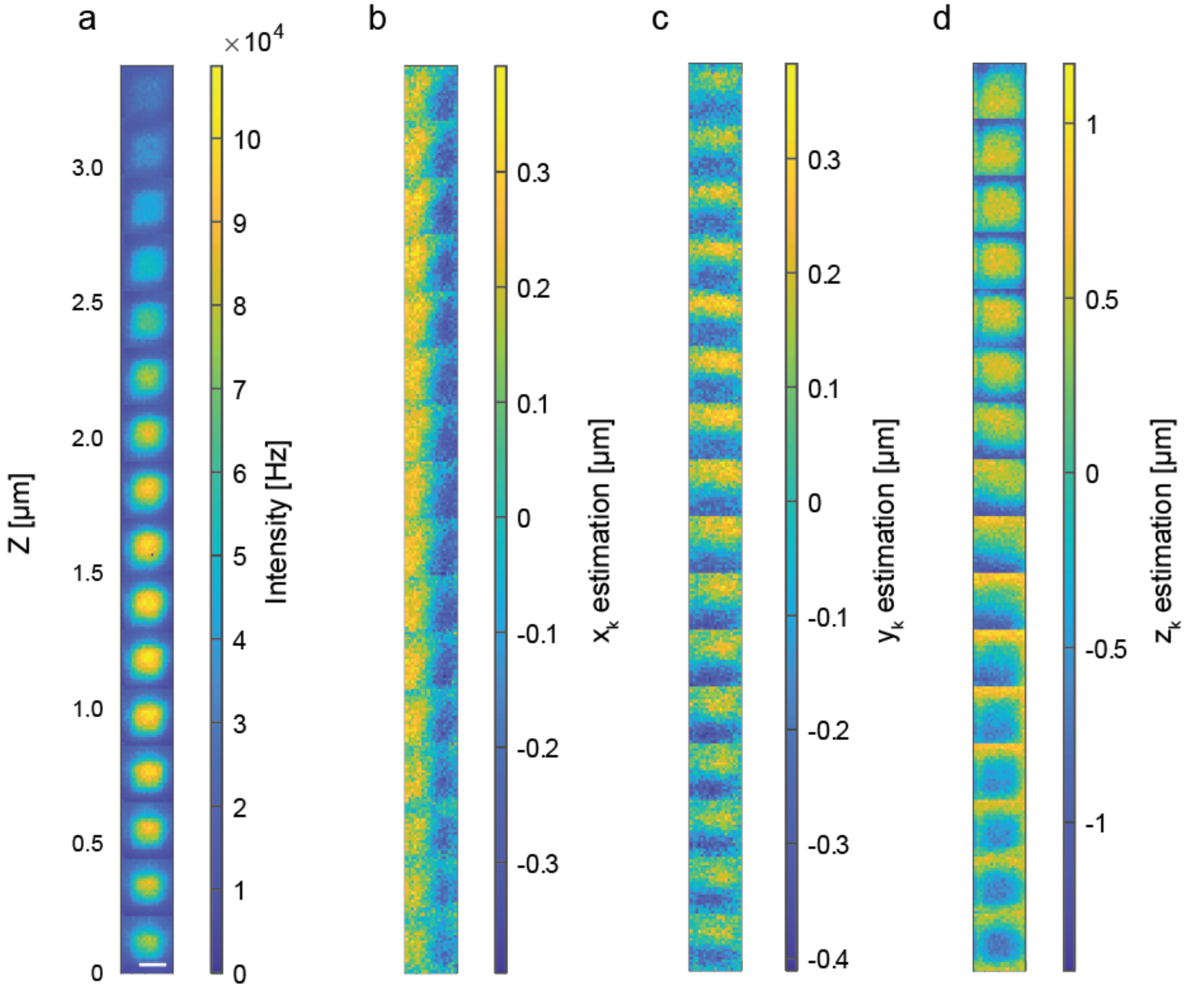

**Figure S3. 3D scan of a 110 nm fluorescent polystyrene nanoparticle.** NP was scanned using the piezoelectric stage with EOD and TAG lens turned on. (a) NP intensity. (b-d) Estimated X (b), Y (c), and Z (d) position of NP during the scan. Scale bar = 1 µm.

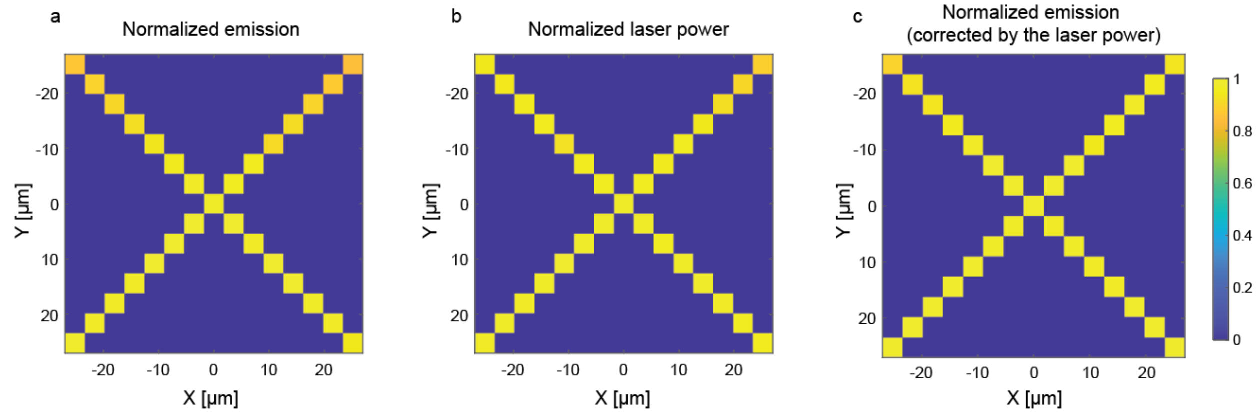

**Figure S4. Galvo scanning range determination.** A slide with uniformly distributed fluorophores was used to establish the galvo scanning range. Yellow scares denote sampled areas, and blue squares are unsampled. (a) Normalized emission at various scan positions, showing a minor drop in collection efficiency at the scan edges. (b) Normalized laser power at different scan positions, showing a minor reduction in laser power at the scan edges. (c) Corrected normalized emissions after correction for the laser power at each scan position. These data suggest a tracking range of at least ± 25.5 μm in both X and Y directions.

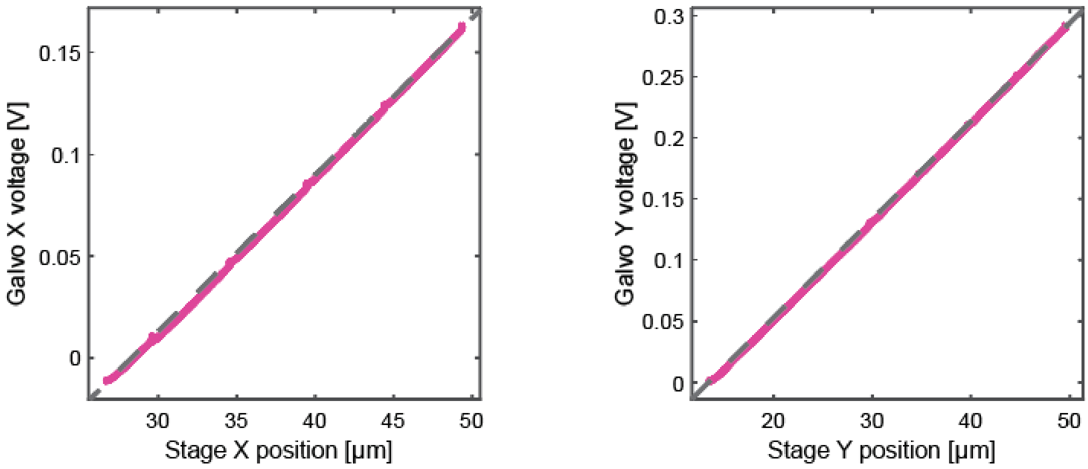

**Figure S5. Galvo mirror calibration.** Galvo positions plotted against stage positions when tracking an immobilized NP moved using the piezoelectric stage. The red lines indicate the experimental results, while the grey dashed lines are the linear fitting results. The fitted slopes of these two plots are 0.0077 and 0.0080 V/μm for X and Y, respectively.

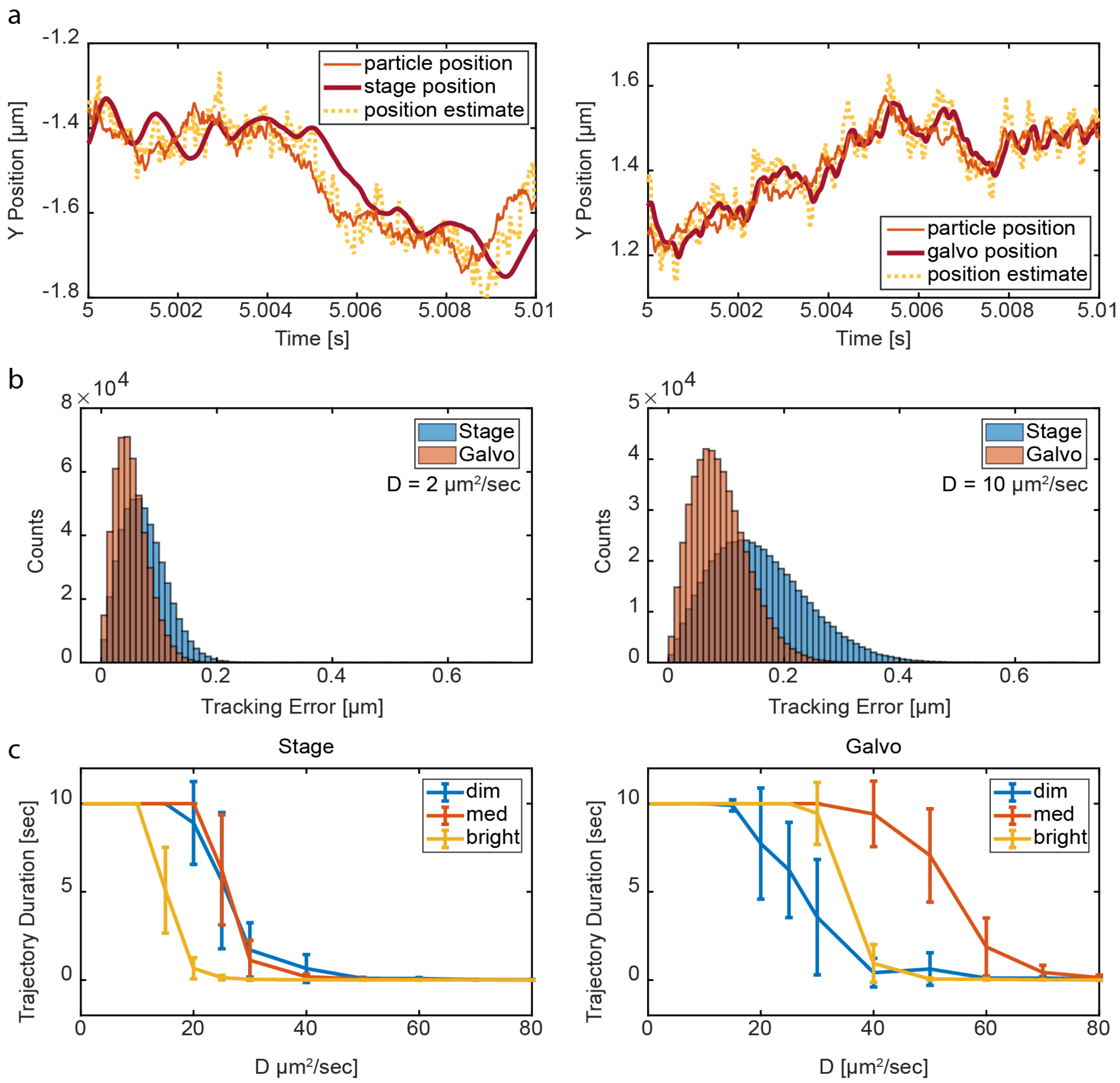

**Figure S6. Simulated galvo and stage tracking performance versus diffusion coefficient.** (a) Example trajectory segment of medium intensity (140 kHz) simulated data along the Y axis for stage (left) galvo (right) tracking. D = 2 μm$^2$/s. (b) Histogram of observed tracking error for each bin of a 10 second trajectory for a slow particle (left, D = 2 μm$^2$/s) and a faster particle (right, D = 10 μm$^2$/s). (c) Duration of particle localization within 0.5 μm of the edge of the laser scan comparing stage tracking (left) and galvo tracking (right). N = 10, max duration = 10 s. The observed brightness of dim, medium, and bright particles are 14 kHz, 140 kHz, and 1401 kHz, respectively. The error bars on the bottom plots are standard deviation values of observed durations.

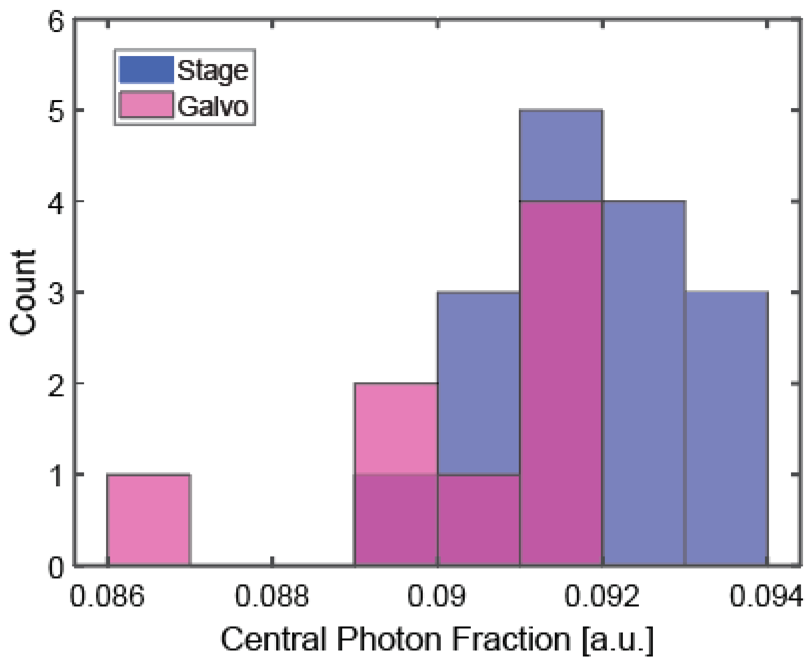

**Figure S7. Histogram of central photon fraction.** The central photon fraction is calculated by tracking immobilized NPs with stage and galvo. The piezo controller was operated in closed-loop for stage tracking, resulting in a central photon fraction of 0.092 ± 0. 001 (N=16). Tracking immobilized NPs using the galvo mirrors resulted in a central photon fraction of 0.091 ± 0.002 (N=8).

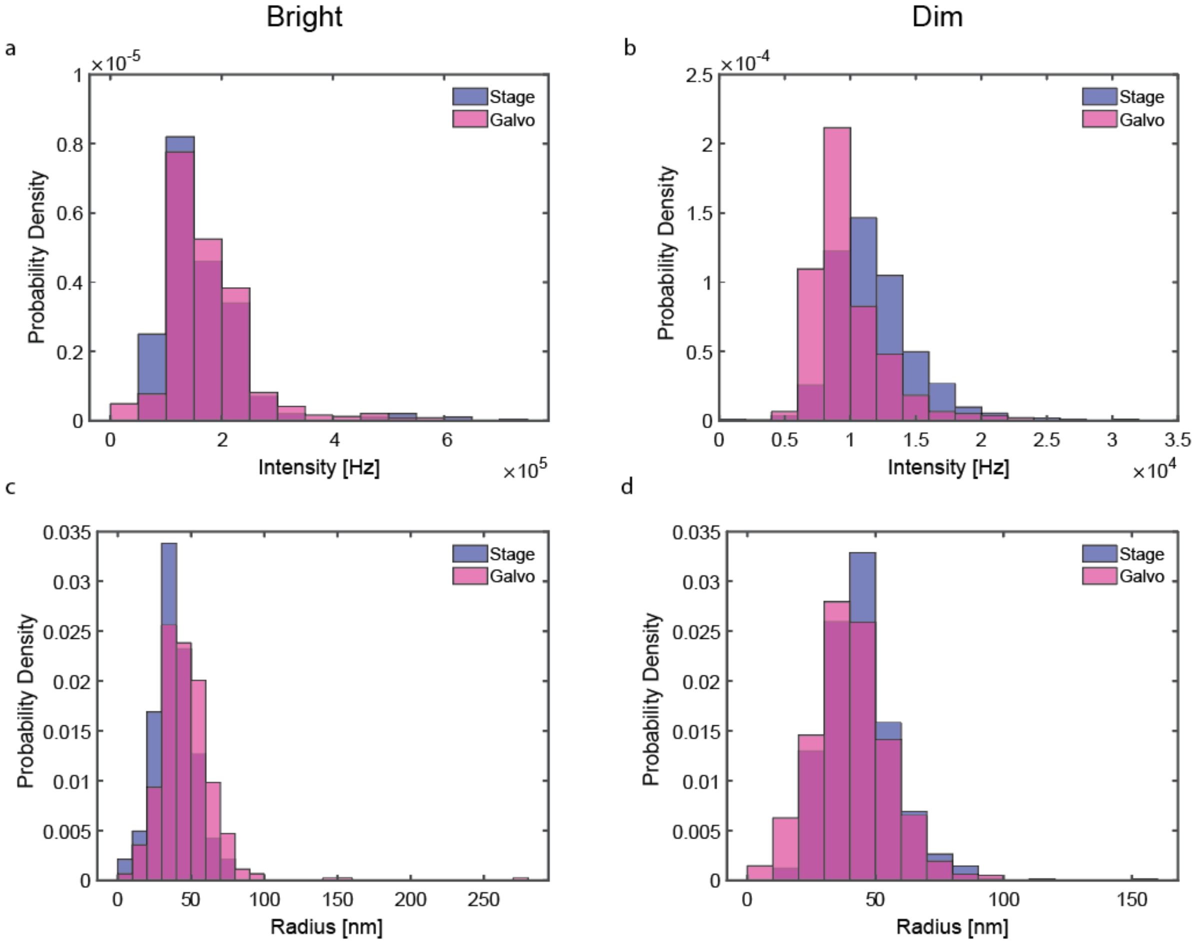

**Figure S8. Tracking bright and dim NPs.** Data from the piezoelectric stage and galvo mirror are shown in blue and pink, respectively. (a) Intensity distributions for bright NPs. Int = 164 ± 83 kHz for stage (n = 283), int = 174 ± 81 kHz for galvo (n = 492). (b) Intensity distributions for dim NPs with intensity less than 12 kHz. Int = 11.7 ± 3.0 kHz for stage (n = 560), int = 9.9 ± 2.9 kHz for galvo (n = 464). (c) Bright NP size distribution, measured by MSD analysis. $R_h$ = 39.9 ± 18.4 nm for stage, $R_h$ = 48.3 ± 21.8 nm for galvo. (d) Dim NP size distribution. $R_h$ = 46.6 ± 15.9 nm for stage, $R_h$ = 43.2 ± 17.9 nm for galvo. Data are expressed as mean ± SD.

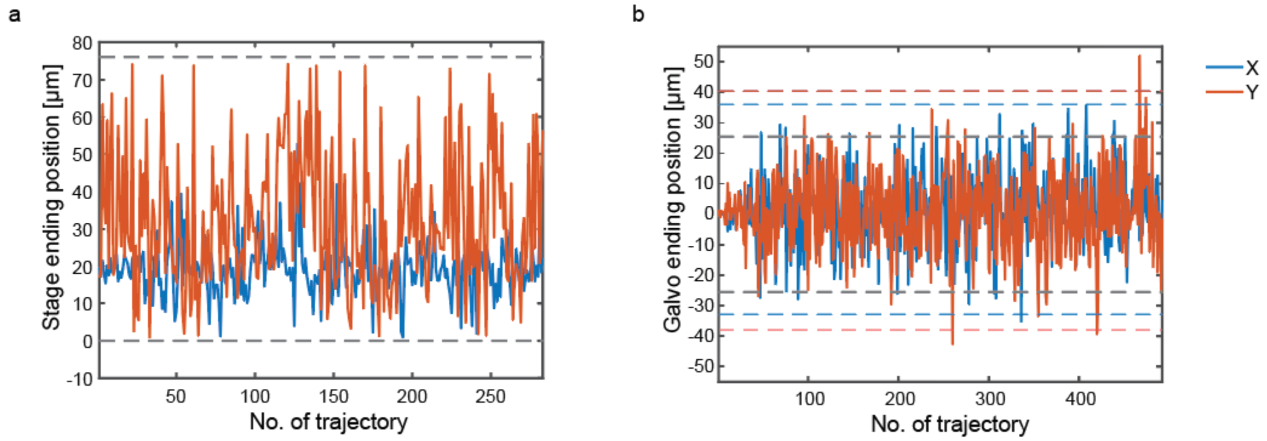

**Figure S9. Ending positions of trajectories for stage and galvo tracking**. (a) Ending positions for stage tracking show a higher tendency of tracking termination due to approaching the limits of stage motion (0-76 µm, grey dashed lines). (b) Galvo tracking ending positions. The grey dashed line shows the ± 25.5 µm tracking boundary with no signal loss. However, the actual tracking range of galvo is larger than ± 25.5 µm (as indicated by the trajectories with ending positions beyond ± 25.5 µm). The galvo range is ultimately limited by the size of lenses and mirrors in the system. The blue (red) dashed lines indicate ending positions three standard deviations away from the mean in the X (Y) direction.

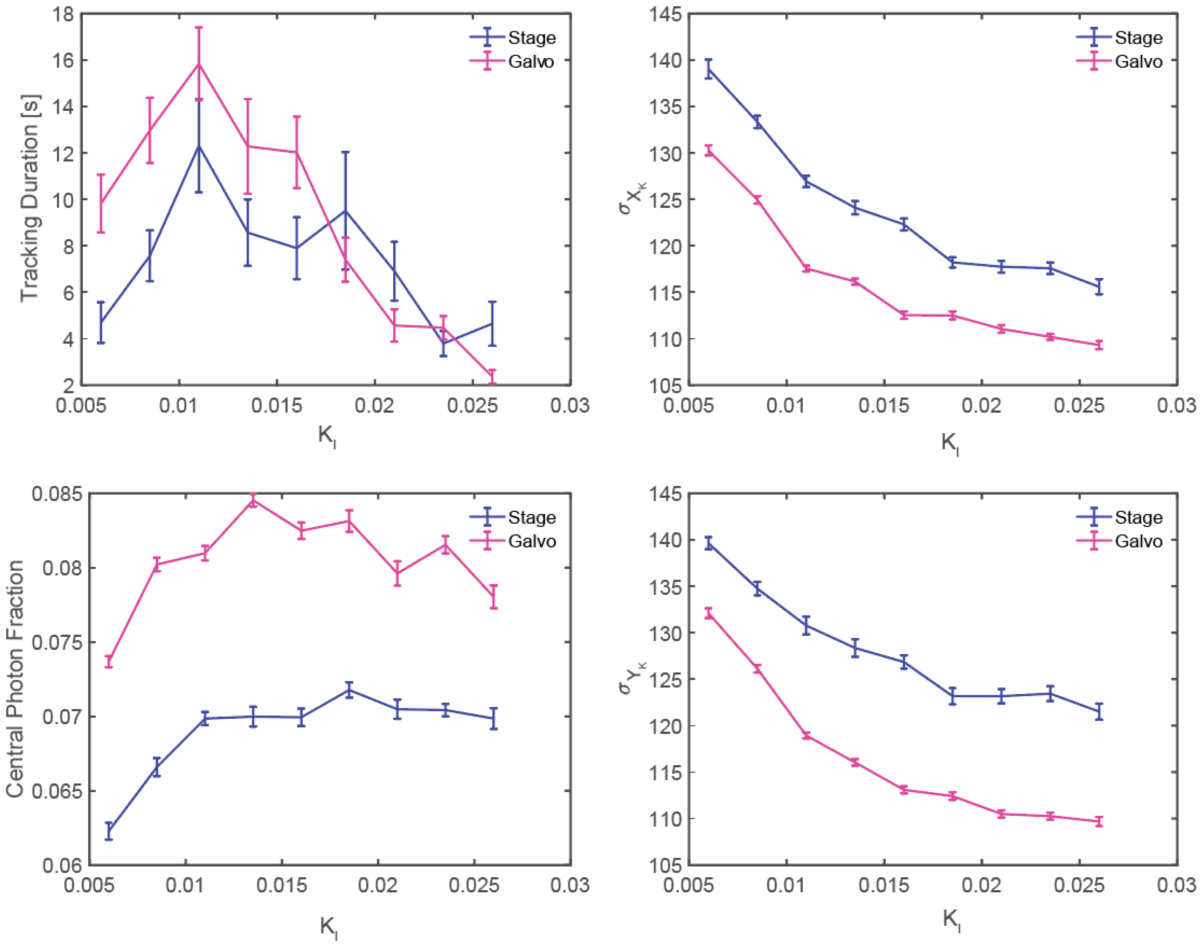

**Figure S10. Tracking performances as a function of $K_I$ for dim NPs.** (a) Tracking duration (b) Central photon fraction. (c) X position estimation precision. (d) Y position estimation precision. NP trajectories with average intensity larger than 12 kHz were removed. The range of $K_I$ is 0.006 to 0.026 with a 0.0025 increment. The error bars indicate the standard error of each parameter measurement.

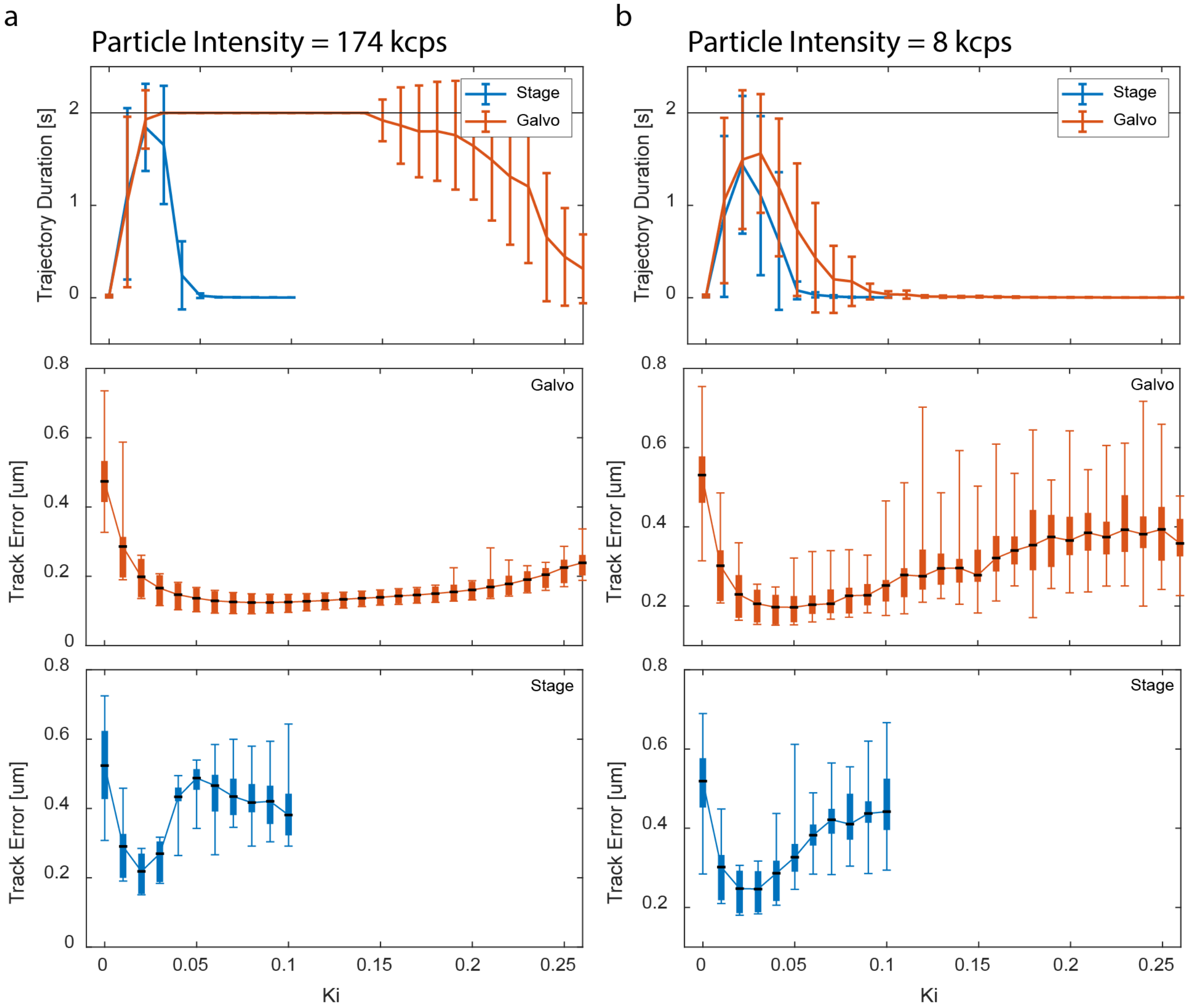

**Figure S11. Simulated tracking of bright and dim particles.** Simulated tracking performance as a function of the $K_I$ value with both bright (174 kHz) and dim (8 kHz) signals to correspond to experimental conditions. Top row: Tracking duration (max = 2 s) for 10 trials each with D = 2, 5, 10, 20, and 30 $\mu m^2/s$ for a total of 50 trials. The error bars are standard deviations. Middle row: mean tracking error for each galvo tracking 2D trajectory prior to escape. Bottom row: Mean tracking error prior to particle escape for piezoelectric stage tracking at each $K_I$.

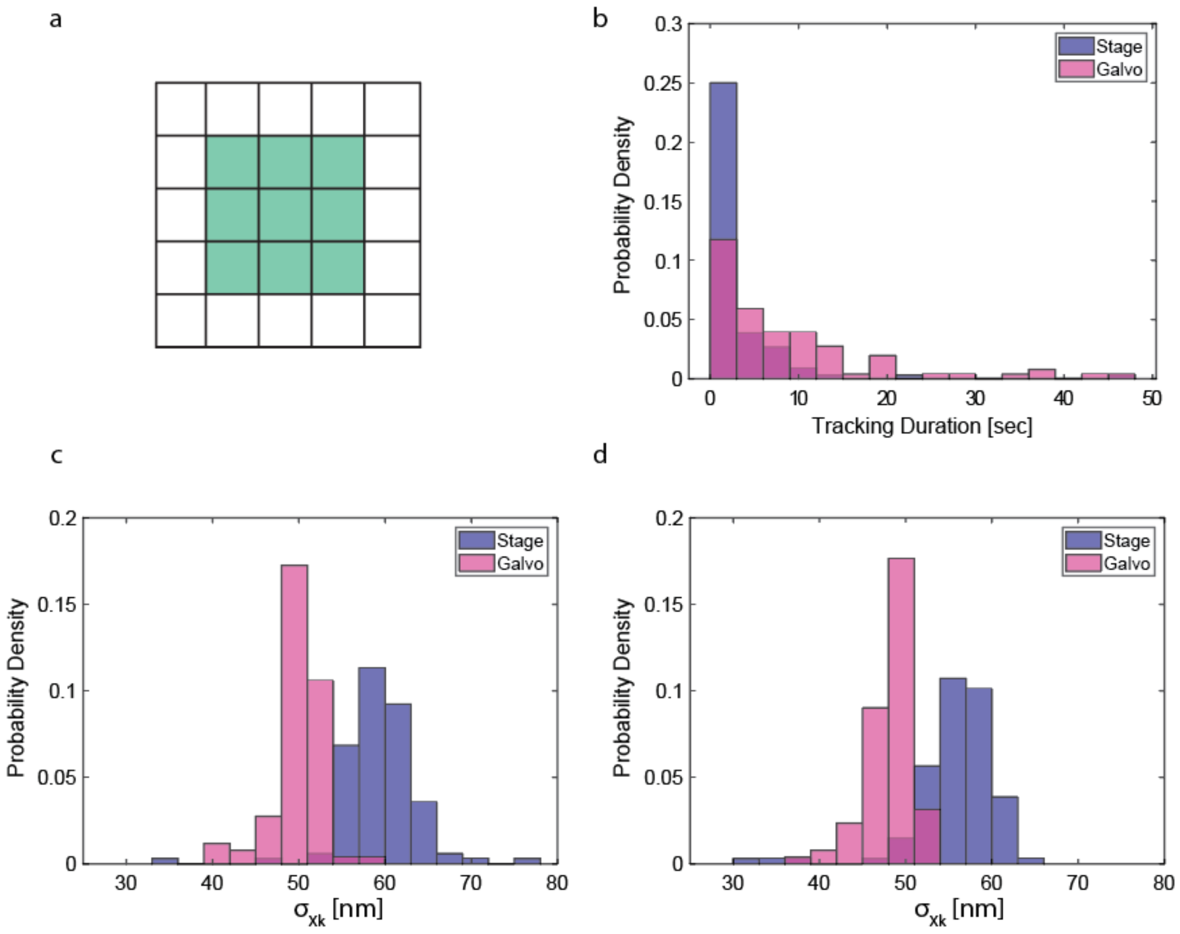

**Figure S12. Stage and galvo tracking performance with a 3 × 3 laser scanning pattern.** (a) 3 × 3 laser scanning pattern. (b) Histograms of tracking duration. Stage: 3.0 ± 5.3 s. Galvo: 8.7 ± 9.8 s. ($p < 0.01$, two-tailed t-test) (c) Histograms of localization precision in X. Stage: 59.3 ± 4.3 nm. Galvo: 49.9 ± 2.7 nm ($p < 0.01$). (d) Histograms of localization precision in Y. Stage: 55.6 ± 4.8 nm. Galvo: 48.4 ± 2.7 nm ($p < 0.01$). Data are expressed as mean ± SD.
